# Connectivity in the human and the monkey brain probes causal involvement of the fornix in Mild Cognitive Impairment and Alzheimer’s Disease

**DOI:** 10.1101/2024.12.02.625865

**Authors:** Vassilis Pelekanos, Shaun Warrington, Elsie Premereur, Jessica de Boer, Stamatios N. Sotiropoulos, Anna S. Mitchell, the Alzheimer’s Disease Neuroimaging Initiative

## Abstract

Alzheimer’s disease (AD) is characterised by memory loss and severe deficits in cognitive function associated with neural degeneration in a network of brain regions. However, little is known about those regions’ connectivity patterns and how that differs from mild cognitive impairment (MCI) or healthy aging. To address that, we used diffusion-weighted MRI to determine connectivity across 11 key memory-related regions and their unique set of connections (connectivity fingerprints) to 14 white matter (WM) tracts. One WM tract particularly important for memory, and attractive target for therapeutic interventions in AD, is the fornix. However, determining fornix-specific contributions to memory deficits or therapeutic benefits is difficult, partly because the fornix carries numerous subcortical and cortical projections. To explore that, we additionally examined MRI-derived connectivity across homologous structures in non-human primates before and after fornix transections. We report several important findings. First, that connectivity between the hippocampus and the anterior thalamus (ATh) is strongly compromised in cognitive decline, as is fornix integrity. We also found strong reductions in the hippocampus-fornix and ATh-fornix connectivity in AD, demonstrating that fingerprint divergence across groups in hippocampal CA1 and ATh can identify differences between people with AD and MCI. In AD, we observed also elevated connectivity between WM tracts and the hippocampus or the ATh, suggesting a compensatory mechanism, which, importantly, depends on a viable fornix. We finally demonstrate that certain thalamic nuclei and hippocampal subfields link through the retrosplenial cortex in both species, highlighting its potential role as an alternative target for interventions in memory disorders.

## Introduction

Alzheimer’s disease (AD), the most common cause of dementia, is an increasing global health challenge characterised by loss of episodic memory (the memory of everyday events) and other severe cognitive problems (1). A characteristic feature of AD is the neuropathological changes within the Papez circuit, the brain’s renowned memory system in the temporal lobe (2, 3), notably, atrophy and neurodegeneration in the hippocampus (4, 5), the anterior thalamus (2, 6–8) and the retrosplenial cortex (RSC) (9–11). White matter (WM) fibers connecting these structures are also affected in AD (for a recent review, see 12). One white matter fiber bundle particularly affected is the fornix, which links the hippocampus with medial diencephalon structures (13, 14).

Mild cognitive impairment (MCI) is a transitional stage between normal cognitive aging and dementia. People with MCI are believed to be at high-risk for developing clinically probable AD (15, 16), and integrity of the fornix has been indicated as a predictive biomarker of progressing from MCI to AD (17, 18). Human and non-human primate studies indicate that fornix damage causes severe memory deficits (19–24), while magnetic resonance imaging (MRI) confirms that the fornix is compromised in AD and in MCI (25–29).

The fornix is a prominent deep brain stimulation (DBS) target in AD (e.g., 30, 31), with several ongoing clinical trials, following the unexpected memory enhancement in a morbid obesity patient undergoing DBS in the hypothalamus/fornix (32). However, the fornix’s appropriateness as an effective DBS target in AD has been challenged and its therapeutic benefits questioned (33–35). While the specifics of any DBS are still to be established, given that the fornix carries numerous subcortical and cortical projections, its facilitative effects likely relate to its interconnected structures, rather than the fornix itself (34, 36, 37).

To better understand the mechanisms underlying cognitive decline in AD, and the modulatory mechanisms driving the memory circuit through fornix stimulation, it is crucial to know how key memory- and fornix-related structures in AD are connected, and perhaps more so, how this connectivity differs in MCI and in healthy ageing. We sought to answer this largely unstudied question in age- and sex-matched groups of people with AD, MCI, and healthy controls. We focused on hippocampal, thalamic and cortical regions strongly implicated in memory, and used recent advances in brain imaging tools (38–40) to delineate these regions’ connectivity fingerprints, that is, describe the connections of each region with key white matter tracts we reconstructed on each brain. Importantly, we investigated whether the fornix plays a causal role in the above aims. Fornix-specific contributions to connectivity degradation are difficult to estimate because, often, forniceal lesions are accompanied with lesions in other structures also involved in memory (23, 41). We filled this gap by using MRI data from macaque monkeys with bilateral fornix transections, and examined connectivity profiles across the same homologous WM tracts following a standardised WM reconstruction pipeline across species (39, 42). Also, we explored region-to-region structural connectivity in both species.

Our results provide important insights for understanding connectivity across gray matter structures and white matter pathways in healthy ageing and how this compares in people with AD or MCI, as well as to macaques with fornix damage. As such, the present study also contributes to our knowledge of the comparative architecture of memory-related neural circuits between humans and macaques alongside helping to identify alternative neuromodulation targets with potential clinical significance for DBS in Alzheimer’s disease.

## Results

### Tract-to-ROI connectivity fingerprints

We examined white matter tracts implicated in memory and higher cognition, homologous across human and macaques, which we reconstructed using cross-species tractography (38, 39). Except for the fornix, which was eventually transected (thus not included) in the monkeys, all tracts were common and successfully generated in each individual brain, each species. We also examined subcortical/cortical regions of interest (ROIs), largely interconnected by the fornix, established correlates of episodic memory, exhibiting well-documented abnormalities in MCI and AD (2, 43, 44). We generated connectivity fingerprints, that is, maps describing the subcortical/cortical terminations of each tract (38).

In both species, we found that all hippocampal subfields were prominently connected to the temporal subsection of the cingulum bundle (CBT), the inferior fronto-occipital fasciculus (IFO), and (less so) the uncinate fasciculus (UF). In humans, all hippocampal connections to the CBT remained unaffected in AD and MCI, but connections to the fornix were strongly degraded in AD, and to a lesser extent in MCI, compared to healthy controls, in both hemispheres (Fig. 1). This indicates that, in the healthy brain, the hippocampus is strongly connected to both the CBT and the fornix, but it is the degradation of the latter pathway only that relates to AD. In fact, compared to controls, AD and MCI groups had slightly stronger connections between the hippocampal subfields and the CBT in each hemisphere, attributing a potentially compensatory role to this pathway. This speculation is supported by the monkey dataset, where, the fornix-transection post-operative MRI scans (post-op) exhibited stronger presence of the CBT tract in all hippocampal subfields’ profiles (except in right CA1) compared to the pre-op. The IFO tract had a notable presence in all hippocampal subfields’ connectivity profiles too, with the AD patients having stronger connection compared to the other two groups. In contrast to the human patients, connectivity between IFO and all hippocampal subfields was greatly reduced in the monkeys post-op, suggesting that a viable fornix is essential for the increased hippocampal-IFO connectivity observed in AD. A similar connectivity pattern was observed in the post-op connections between the left hemisphere’s CA1, CA3 and CA4 and the UF tract.

**Fig. 1.**
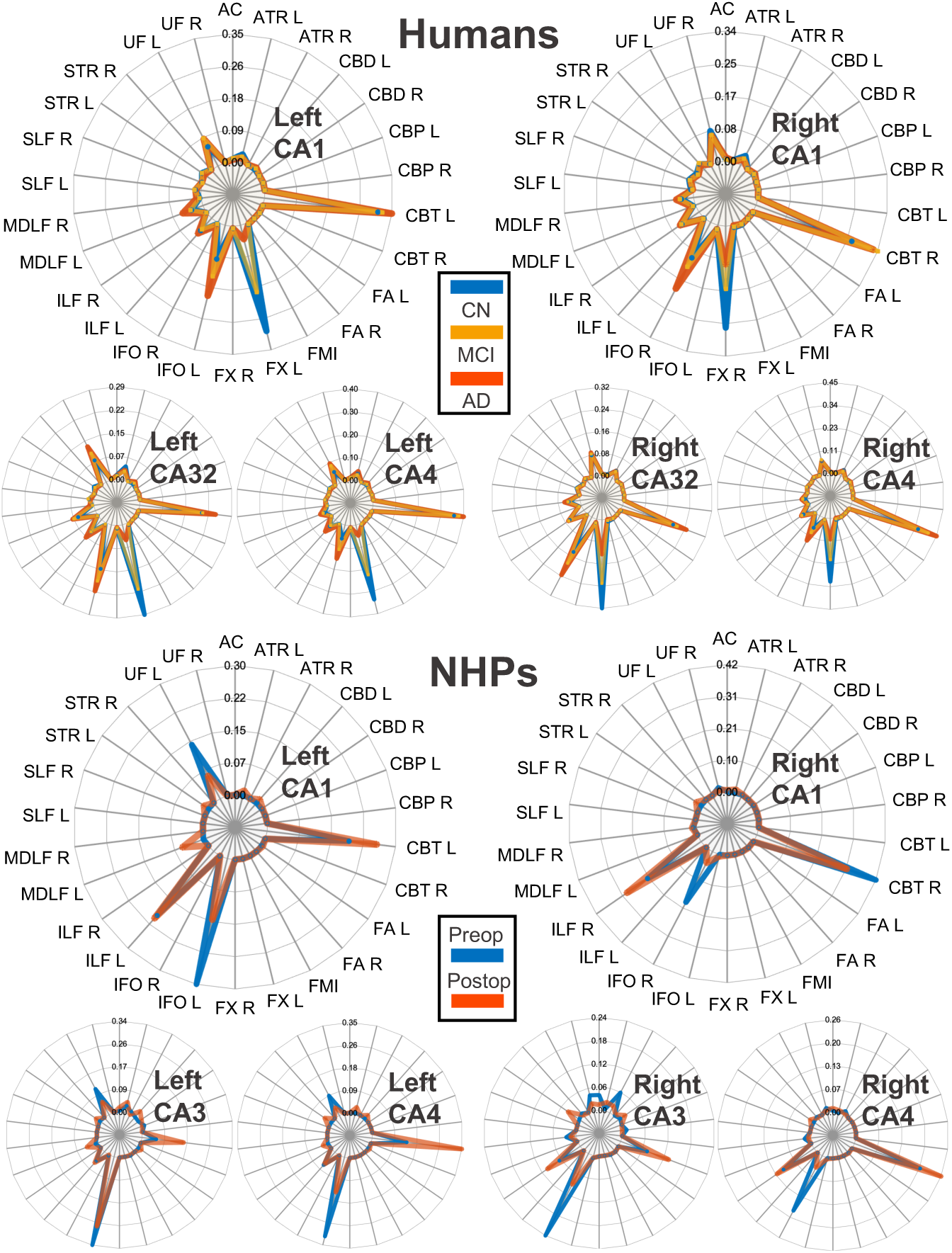
Connectivity fingerprints between hippocampal subfields and white matter tracts, in the left and right hemisphere. Top panel shows human data for healthy controls (CN), mild cognitive impairment (MCI) and Alzheimer’s disease (AD) patients. Bottom panel shows the non-human primate (NHP) data before (pre-op) and after (post-op) the fornix transection operations. Given that the fornix was transected in the post-op group, connectivity fingerprints to the fornix were not calculated. Tracts abbreviations: fx: Fornix; atr: Anterior Thalamic Radiation; cbd: Cingulum subsection dorsal; cbp: Cingulum subsection: perigenual; cbt: Cingulum subsection: temporal; fa: Frontal Aslant; ifo: Inferior Fronto-Occipital Fasciculus; ilf: Inferior Longitudinal Fasciculus; mdlf: Middle Longitudinal Fasciculus; slf1:Superior Longitudinal Fasciculus 1; str: Superior Thalamic Radiation; uf: Uncinate Fasciculus; ac: Anterior Commissure; fmi: Forceps Minor. L: left hemisphere; R: right hemisphere.

Within each species, the subicular complex regions (**Supplementary Fig. 1**) exhibited almost identical connectivity profiles, that were like the connectivity profiles of CA3 or CA4 hippocampal subfields. In monkeys, the subicular complex, like the hippocampus, was prominently connected to the CBT and IFO tracts. In humans, the subicular complex was connected strongly to the CBT (and, like the hippocampus, it remained unaffected in AD and MCI), and to a lesser extent to the fornix where connectivity was considerably reduced in AD compared to MCI and control groups.

The anterior thalamic nucleus (ATh) connection to the fornix was also strongly compromised in AD, compared to MCI and control groups that exhibited similar strengths (Fig. 2). However, connectivity between the ATh and the anterior thalamic radiation (ATR) tract was increased in AD patients compared to people with MCI and healthy controls. In our monkey dataset, in contrast, connectivity between the ATR tract and all thalamic nuclei (except for the right anteroventral nucleus; AVN) was degraded by fornix transection compared to pre-op. Like the cross-species hippocampal subfields results, these human and monkey findings together suggest that a viable fornix is essential for the increased thalamic-ATR connectivity observed in AD.

**Fig. 2.**
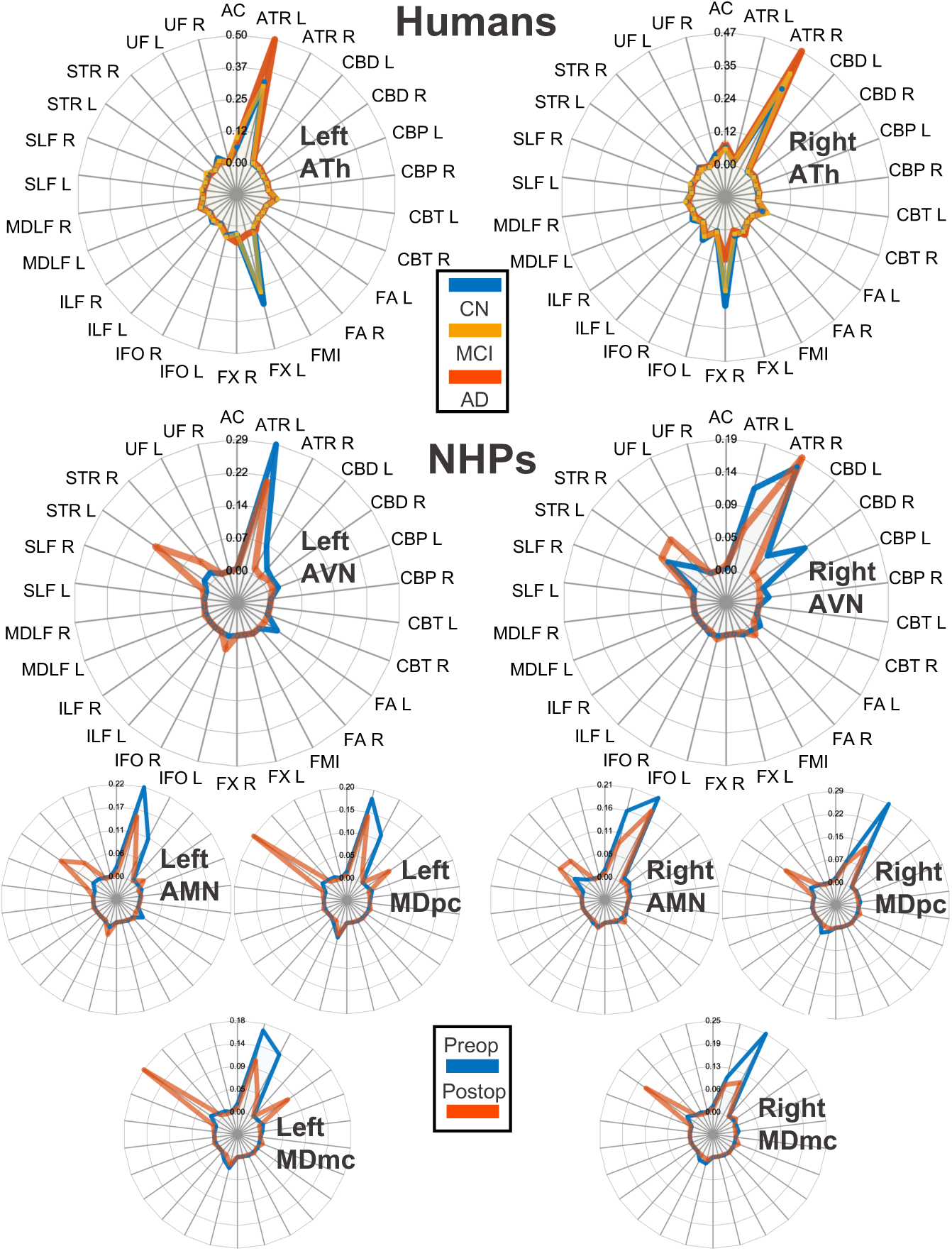
Connectivity fingerprints between thalamic nuclei and white matter tracts in the left and right hemisphere. Top panel shows human data for healthy controls (CN), mild cognitive impairment (MCI) and Alzheimer’s disease (AD) patients. Bottom panel shows the non-human primate (NHP) data before (pre-op) and after (post-op) the fornix transection operations.

Strong connectivity to the ATR tract was found for the remaining thalamic nuclei we examined in both humans and macaques. In humans, the MDl and MDm thalamus had almost identical connectivity profiles (with the MDl being additionally connected to the Superior Thalamic Radiation; STR) across the AD, MCI, and control groups in each hemisphere (**Supplementary Fig. 1**).

Connectivity patterns of the retrosplenial cortex (RSC) are shown in Fig. 3. The dorsal subsection of the cingulum bundle (CBD) was predominantly present in the connectivity profiles of RSC area 29 in both species, while, in monkeys, area 29 showed almost identical connectivity profile to neighbouring RSC area 30, in both hemispheres. Interestingly, monkey areas 29 and 30 were connected to the IFO tract too, especially post-op, and human area 30 showed increased connectivity to the CBT tract.

**Fig. 3.**
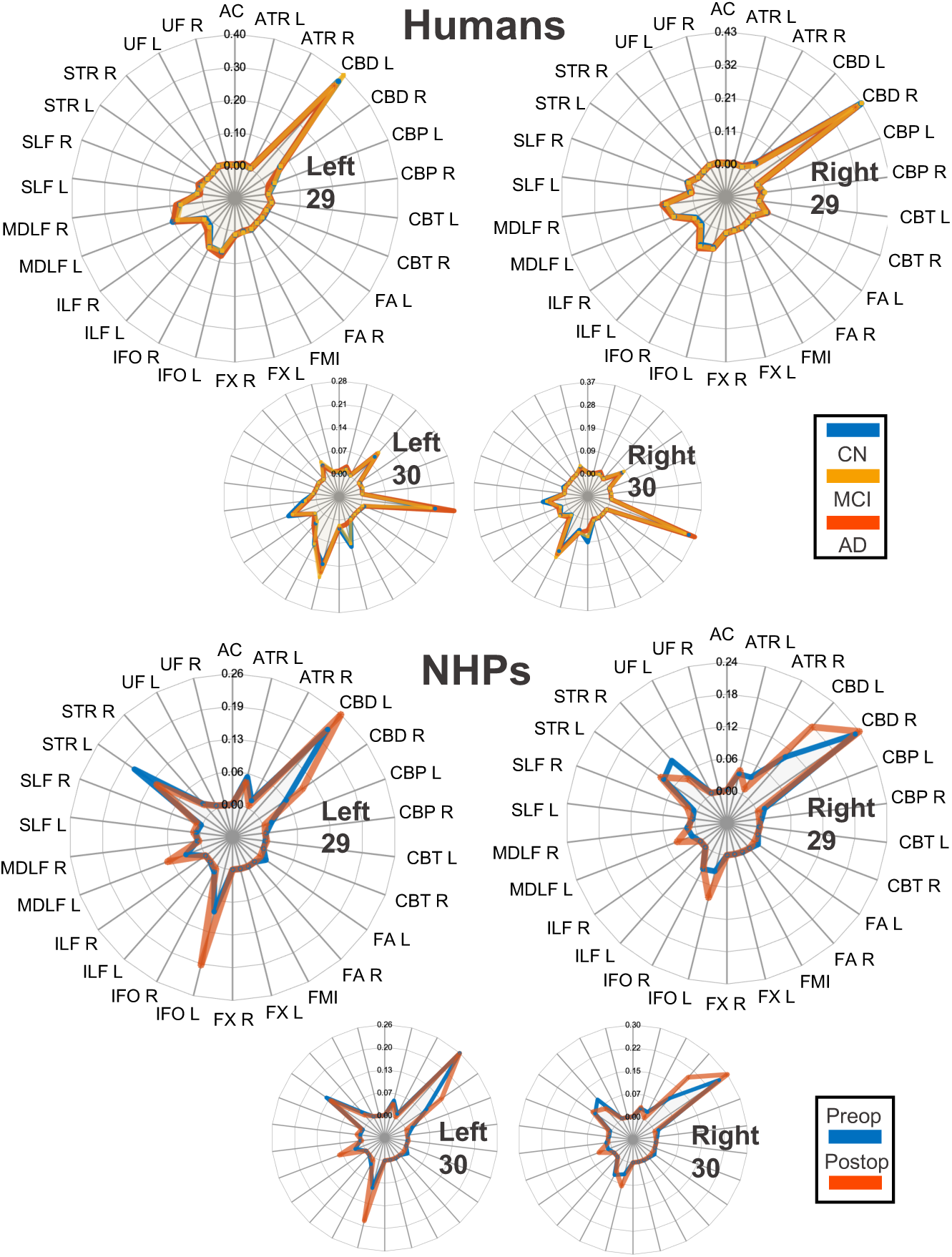
Connectivity fingerprints between the retrosplenial cortex ROIs and white matter tracts, in the left and right hemisphere. Top panel shows human data for healthy controls (CN), mild cognitive impairment (MCI) and Alzheimer’s disease (AD) patients. Bottom panel shows the non-human primate (NHP) data before (pre-op) and after (post-op) the fornix transection operations.

### Divergence of connectivity fingerprints across groups

We compared connectivity fingerprints across groups, to address how a given tract-to-ROI connectivity fingerprint differs between groups, using the metric of Kullback-Leibler (KL) divergence (see *Materials and Methods*). We focused on the fingerprints of hippocampal CA1 and anterior thalamus (ATh) in both hemispheres, whose voxels’ KL divergence distributions are shown in Fig. 4A and Fig. 4B respectively. Comparing CA1’s divergence between [Alzheimer’s Disease (AD) vs healthy controls] and [mild cognitive impairment (MCI) vs healthy controls], we notice that, while distributions have the same maximum values (∼4.5), distributions are less right-skewed in the [AD vs controls] test, with higher proportion of CA1 voxels having high divergence values, than in the [MCI vs controls]. Wilcoxon rank sum tests confirmed that the greater divergence in [AD vs controls] was statistically significant in both the left [*z* = 32.2, *p* <.001] and the right [*z* = 14.3, *p* <.001] hippocampal CA1. This indicates that the connectivity patterns of CA1 were more divergent in the AD brain compared to healthy controls, than in MCI compared to controls, suggesting that the overall tracts-to-ROI connectivity in CA1 can better identify differences between people with AD and people with MCI. Divergence distributions in the ATh (Fig. 4B) also suggest a higher proportion of ATh voxels with high divergence values in the [AD vs controls] than in the [MCI vs controls] test. This was confirmed by statistically significant Wilcoxon rank sum tests in both the left [*z* = 16.2, *p* <.001] and the right [*z* = 6.5, *p* <.001] ATh, indicating that connectivity patterns within the ATh can also identify differences between people with AD and people with MCI.

**Fig. 4.**
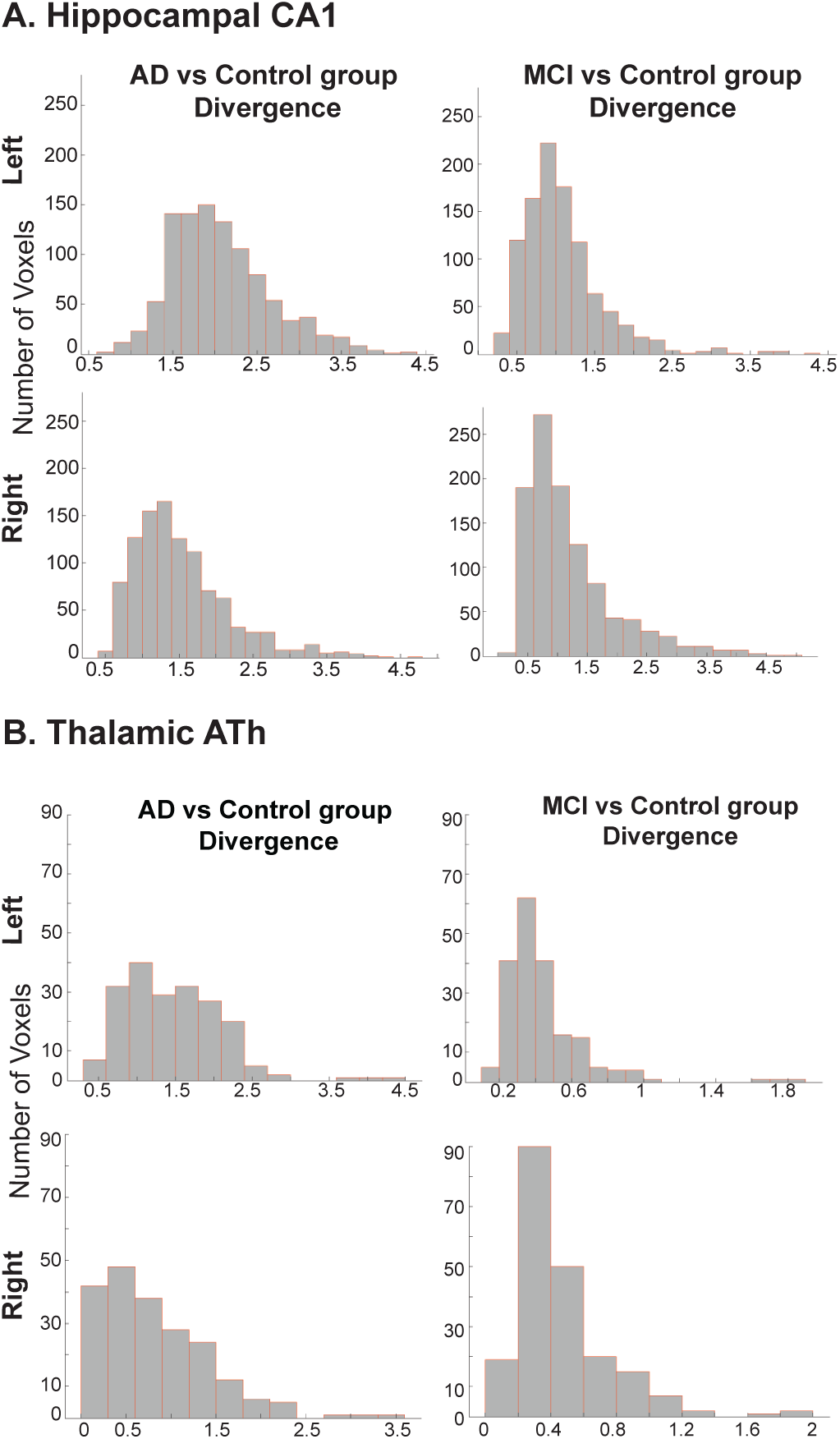
Hippocampal CA1 subfield (**A**) and anterior thalamus (ATh) (**B**) connectivity fingerprint KL divergence between Alzheimer’s Disease (AD) and healthy controls (left column), and between mild cognitive impairment (MCI) and healthy controls (right column). Data for the left and right hemispheres shown in top and bottom panels respectively. Y axes show the number of voxels. X axes show a given group’s divergence to the average connectivity fingerprint of the given ROI in the other group.

### White matter tracts microstructural integrity

We have used Fractional Anisotropy (FA) as a proxy for the white matter tracts’ microstructural integrity. FA is a diffusion MRI metric for water diffusion asymmetry (anisotropy) within a voxel (e.g., 45), with higher FA values representing increased anisotropy, i.e., increased directionality of diffusion, reflecting higher microstructural integrity and fiber organisation. FA is estimated by the Diffusion Tensor Imaging model, which is limited in modelling crossing-fibers (e.g., 46), although recent studies suggest that it is a reliable proxy for white matter axons integrity (47, 48).

We found that, for the majority of our tracts of interest, FA was reduced in AD, compared to MCI and controls in humans, and in fornix-transected (post-op), compared to intact (pre-op), monkeys (Fig. 5). Exceptions were observed in the human peri-genual subsection of the cingulum (CBP), first branch of the superior longitudinal fasciculus (SLF1), and superior thalamic radiation (STR) tracts where FA was slightly higher in AD than MCI, and in the inferior fronto-occipital (IFO) tract where MCI and AD scored higher than the control group. This latter result accords with our finding of increased hippocampal-IFO connectivity in AD and MCI compared to controls (Fig. 1), providing support to this tract’s potential compensatory role in cognitive decline (see *Discussion*). In the monkeys, the CBP, inferior longitudinal fasciculus (ILF), and the uncinate fasciculus (UF) scored higher post-op.

**Fig. 5.**
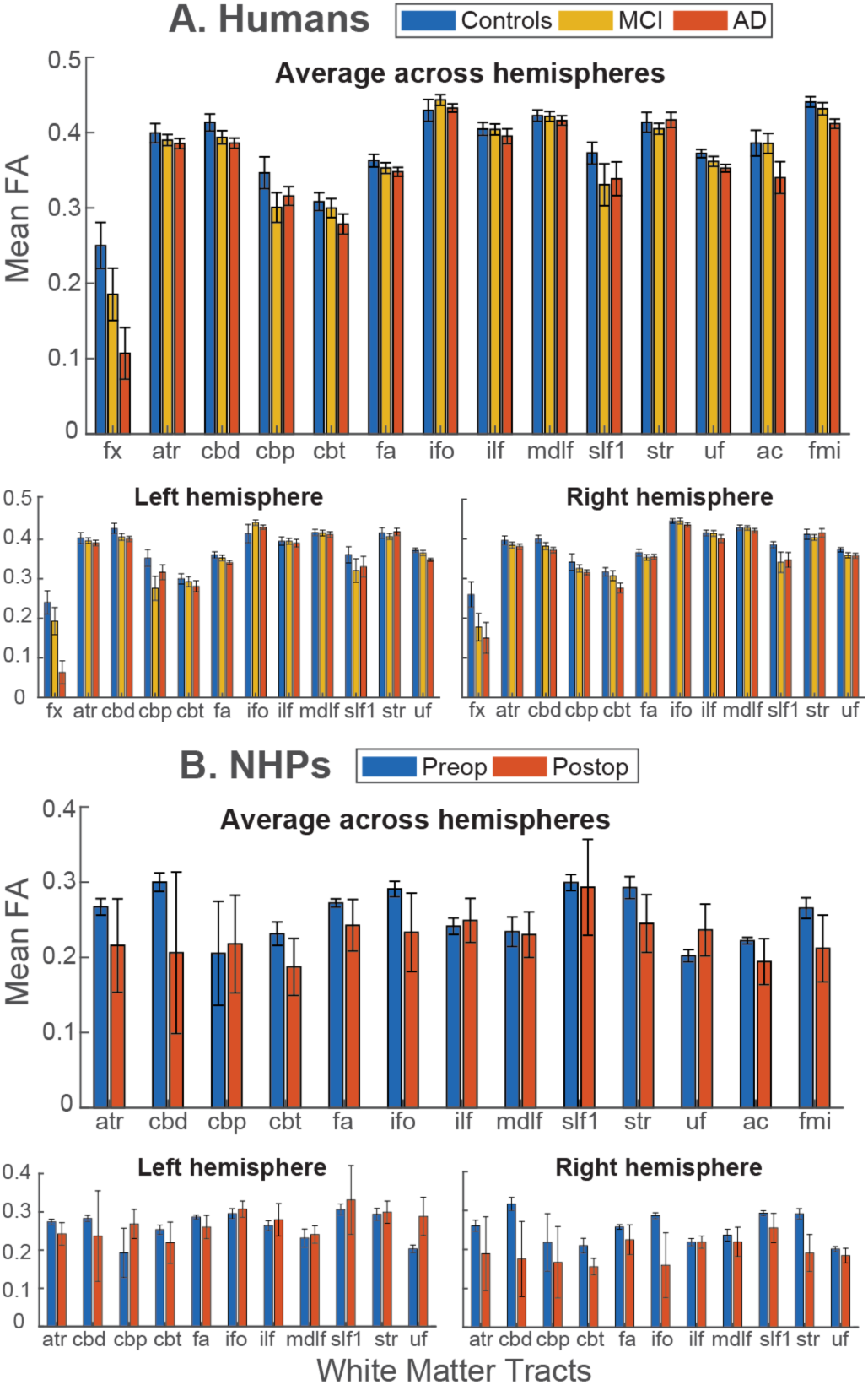
Group mean Fractional Anisotropy (FA) across white matter tracts of interest, in Alzheimer’s Disease (AD), mild cognitive impairment (MCI) and healthy controls (**A**), and non-human primates (NHPs) preoperative (pre-op) and postoperative (post-op) fornix transection (**B**). Top panels show FA values averaged across hemispheres within each tract, and bottom panels show FA in each hemisphere separately. Note that AC and FMI are commissural tracts, thus, each was reconstructed as a whole. Due to the group’s fornix transection post-op, the fornix is excluded from NHPs’ data in panel B. Error bars indicate standard error of the means. See Fig. 1 for tracts abbreviations.

Importantly, this analysis revealed that fornix FA was strongly compromised in humans with AD, providing further support to previous studies (see *Introduction*). The AD group showed the lowest mean FA values for the fornix, followed by MCI compared to controls. This effect of group was highly significant [*χ*^2^(2) = 11.87, *p* = 0.003]. Post-hoc comparisons showed a significant difference to the control group for AD [t(57)= 3.52, *p* = 0.0017] but not for MCI [t(57) = 1.61, *p* = 0.1124]. The remaining bilateral tracts showed similar trends, with all except the IFO showing the highest mean FA in the control group. However, the effect varied between tracts, with notable differences in FA between the control and AD group for the CBP, CBT and SLF1, but not the IFO, ILF, and MDLF tracts. This was reflected in a significant interaction of group with tract [*χ*^2^(22) = 59.4, *p* = 0.00003] when the bilateral tracts (excluding the fornix) were modelled jointly. Importantly, there was no interaction between group and hemisphere, so that the FA values could be averaged across hemispheres and combined with the two commissural tracts (AC and FMI) in a joint model analysis. This showed a main effect of group that approached significance [*χ*^2^(2) = 5.52, *p* = 0.06316], with post-hoc comparisons to the control group showing a significant difference for the AD [t(57) = 3.52, p = 0.0017] but not for the MCI group [t(57) = 1.61, p = 0.1124]. When the models were repeated after removing data points identified as outliers according to residual analysis (see *Materials and Methods*), the effects of group retained the reported trends with group, but with stronger effect sizes, confirming that the effects were not biased by outliers.

Interestingly, FA at the fornix was reduced also in the healthy control group compared to all other tracts we examined. Given that all human participants in our study are of older age (see *Materials and Methods*), this observation suggests that the fornix is more vulnerable to the well-known age-related degradation of white matter integrity (49–51). This accords with previous reports of reduced FA in the fornix in older adults (52–54). Emphatically, it has been also shown (55) that hippocampal atrophy in healthy aging is specifically associated with reduced FA in the fornix, but not the cingulum bundle.

### ROI-to-ROI connectivity

Our results so far indicate that fornix microstructural integrity and, importantly, its connectivity to the hippocampus and the anterior thalamus (ATh) are strongly degraded in AD, and to a lesser extent, in MCI. Recently, Aggleton et al. (56) suggested that the relationship between the hippocampus and the ATh is not hierarchical. Rather, the ATh may provide a route to the hippocampal formation indirectly, with candidate cortical sites including the retrosplenial cortex (RSC; areas 29, 30). To determine how these regions are connected to one another in our human and monkey cohorts, we conducted a secondary analysis, exploring structural connectivity ROI-by-ROI. We quantified differences in connectivity across groups, using the number of streamlines connecting a seed and a target ROI, in the context of MRI-based probabilistic diffusion tractography (see *Materials and Methods*).

Examining differences between AD patients and healthy controls (Fig. 6A) revealed that the AD group had significantly reduced structural connectivity between the hippocampal system and the thalamic nuclei, bilaterally, as well as within the hippocampus. Strongest differences were found in the left hemisphere, between CA1 and ATh, between CA32 and ATh, between CA4 and ATh, between the subiculum and ATh, and between CA32 and the mediodorsal thalamus (MDl). Significantly reduced connectivity for the AD patients was also observed between the RSC and the thalamus (RSC area 30 to ATh, and area 29 to MDl). We then examined connectivity differences between people with MCI and healthy controls (Fig. 6B). Differences were largely found in almost identical ROI pairs as in the AD-controls test, albeit less strong or statistically non-significant. Of particular interest is the significant reductions, for the MCI, between left CA32/CA4/subiculum/presubiculum and left ATh, just like in AD-controls, indicating that these connections are affected early on in cognitive decline. Comparing connectivity differences between AD and MCI groups directly (Fig. 6C) revealed that the AD patients had significantly reduced connectivity in the left hemisphere, between CA32 and ATh, between CA4 and presubiculum, and between subiculum and presubiculum, suggesting these connections are disrupted not before AD clinical manifestation.

**Fig. 6.**
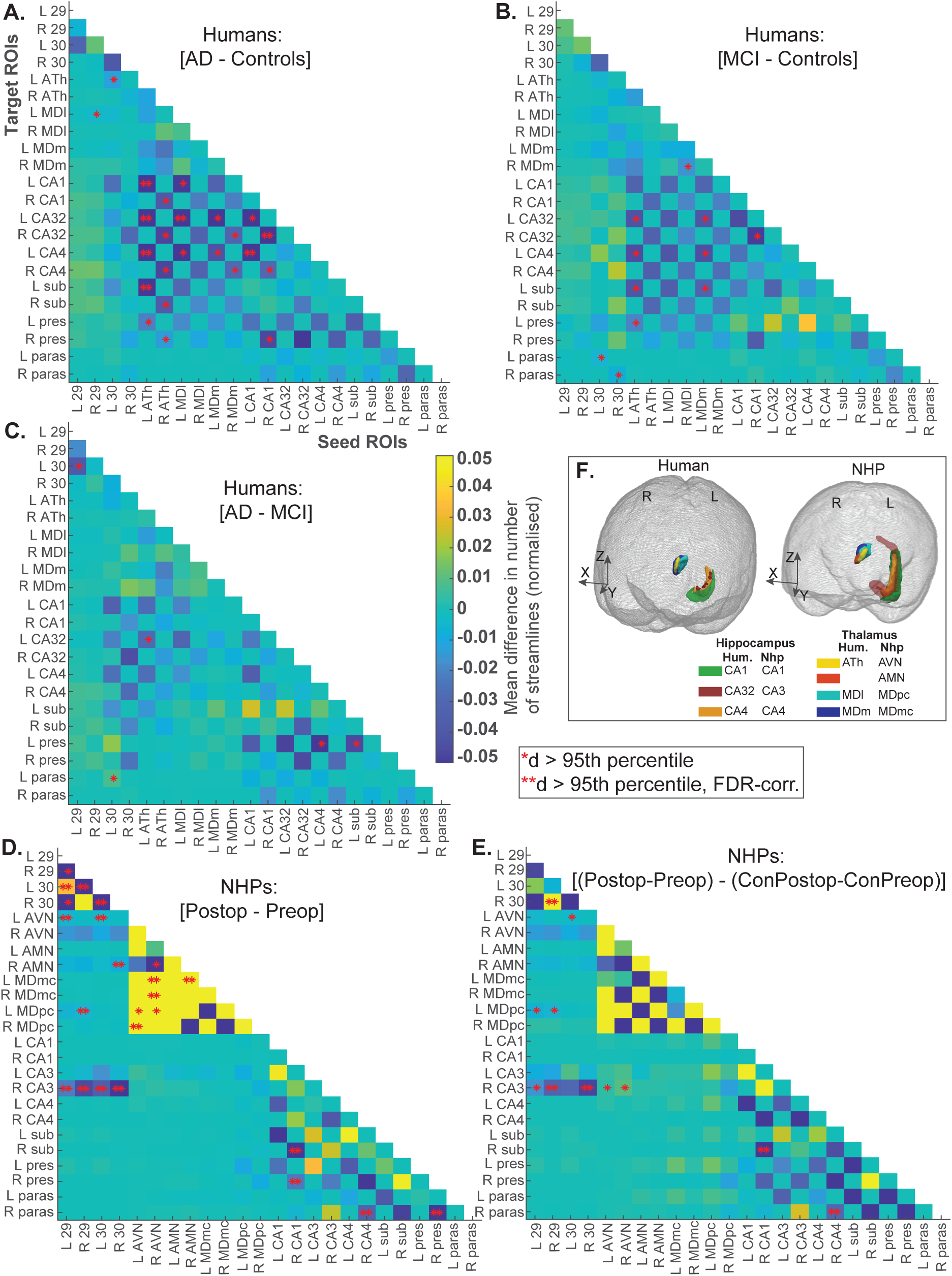
Mean difference in the normalised number of streamlines between Alzheimer’s disease (AD) patients and age- and sex-matched healthy controls (**A**). Values >0 indicate increases in connectivity and values <0 indicate decreases in connectivity for the AD compared to the control group. One asterisk indicates ROI-ROI pairs for which connectivity difference (d) between two given groups was larger than the 95th percentile of a null distribution; two asterisks indicate the ROI-ROI pairs for which d was larger the 95th percentile, FDR-corrected (see *Materials and Methods*). Similarly, comparisons between people with MCI and healthy controls, between AD and MCI, between post-op and pre-op fornix transection non-human primates (NHP), and between the fornix and neurologically intact NHP groups, are shown in **panels B, C, D,** and **E,** respectively. **Panel F** shows core subcortical regions, i.e., the hippocampal subfields and thalamic nuclei, in 3D glass brain representations for the left hemisphere of humans and monkeys in standard Montreal Neurological Institute (MNI) and F99 space, respectively. ROIs abbreviations: Humans: ATh: anterior thalamus (includes anteroventral, anterior medial, and anterior dorsal nuclei; see (84)); MDl: mediodorsal thalamus lateral (parvocellular); MDm: mediodorsal thalamus medial (magnocellular). NHPs: AVN: anteroventral nucleus of the thalamus; AMN: anteromedial nucleus of the thalamus; MDmc: mediodorsal nucleus of the thalamus (magnocellular); MDpc: mediodorsal nucleus of the thalamus (parvocellular). Both species: 29, 30: retrosplenial cortex; CA1/CA3/CA4: hippocampal subfields (in humans, CA3 includes CA2; see (85)); sub: subiculum; pres: presubiculum; paras: parasubiculum. L: left hemisphere; R: right hemisphere.

We also investigated structural connectivity differences post-operatively (post-op) bilateral fornix transection, compared to our animals’ pre-op MRI sessions (Fig. 6D). We found strong fornix-transection-related decreases in connectivity between RSC 29/30 and hippocampal CA3, and (like in the human patients) between the RSC and the anterior and MD thalamus. These results support previous studies that identify the RSC as a key site in learning and memory given its interconnectedness with thalamic and hippocampal structures (9–11, 56, 57), and further indicate that the RSC-CA3 connection is reliant on the relay of neural signals conveyed via a viable fornix. We also found post-op connectivity decreases in the right hemisphere, between the subiculum/presubiculum and CA1, and within the RSC, whereas, intriguingly, fornix-transected monkeys showed significant connectivity increases within the thalamus.

Finally, we analysed an additional dataset from an independent group of monkeys, operated to receive a head-post implanted onto the skull, but remained neurologically intact (see *Materials and Methods*). Therefore, this group was a suitable baseline control (additional to the fornix-transected group’s own pre-op sessions) for our fornix-transected animals. We examined the two groups’ interaction, i.e., the difference [fornix group – control group], that is, [(Post-op - Pre-op) – (ConPost-op - ConPre-op)] (Fig. 6E). This additional analysis showed almost identical connectivity differences to those observed in the fornix group’s [Post-op – Pre-op] test in Fig. 6D, providing support that they are fornix-damage-specific.

### Divergence of connectivity fingerprints across species

Most of our knowledge on the neuroanatomical connections of the retrosplenial cortex (RSC) with the hippocampus and thalamus arises from tracing studies in monkeys and other invasive approaches in rodents and rabbits (9–11, 57, 58). Our monkey and human ROI-to-ROI connectivity results (Fig. 6) accord with these reports, by showing that RSC connectivity to the thalamus and to hippocampal CA3 degraded strongly in fornix-transected monkeys. In humans with AD, RSC-hippocampal reductions did not reach statistical significance, but the RSC-thalamic connectivity was significantly reduced.

To explore how the RSC’s white matter connectivity profiles also compare between monkeys and humans, we calculated KL divergence between a given voxel in macaque RSC to the average connectivity fingerprint of RSC in humans. Divergence maps were represented in the macaque brain space (see *Materials and Methods*), and divergence distributions for areas 29 and 30 are shown in Fig. 7. Connectivity fingerprints in the RSC had a lesser proportion of voxels with high divergence in the [human AD vs monkey post-op] than the [AD vs monkey pre-op] test. Wilcoxon rank sum tests indicated that those differences in divergence were highly significant in both the left [*z* = -27.6, *p* <.001] and the right [*z* = -17.1, *p* <.001] RSC 29, as well as in the left [*z* = -20.3, *p* <.001] and the right [*z* = 12.8, *p* <.001] RSC 30. This indicates that the overall tracts-to-ROI connectivity profile in the RSC is more conserved between the AD neuropathological brain and the monkey fornix-damaged brain, than it is between the AD brain and the monkey intact brain.

**Fig. 7.**
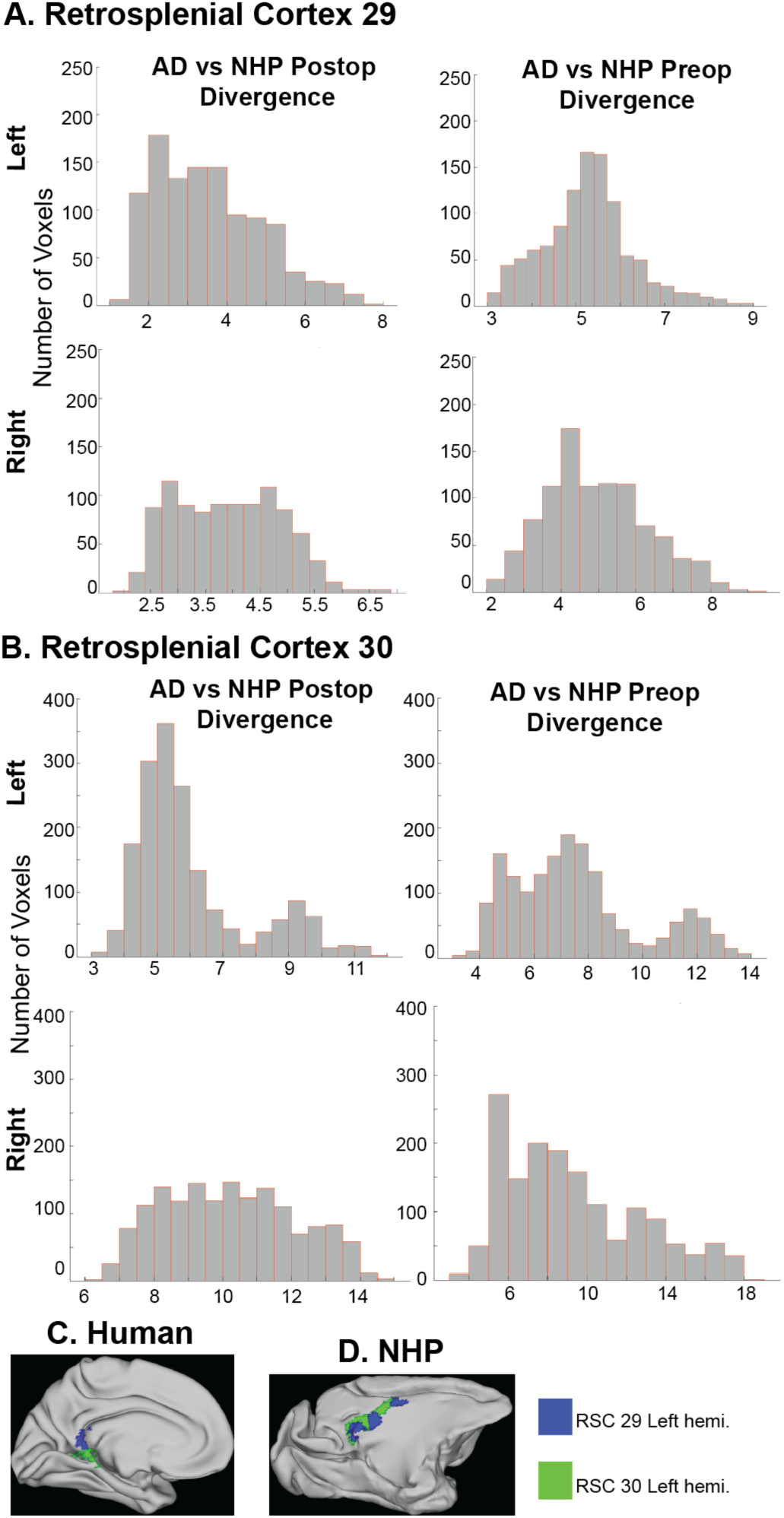
Retrosplenial area 29 (**A**) and 30 (**B**) connectivity fingerprint KL divergence between Alzheimer’s disease (AD) and the macaque monkey fornix post-operative group (left column), and AD and monkey pre-operative group (right column). Data for the left and right hemispheres shown in top and bottom panels respectively. Y axes show the number of voxels. X axes show divergence values between humans with AD to the average connectivity fingerprint of the given ROI in the corresponding monkey group (post-op / pre-op). Panels **C-D** show RSC areas 29 and 30 on mid thickness surface representations in MNI and F99 standard space for the human and monkey respectively.

## Discussion

To evaluate how higher cognition-related white matter tracts drive the connections of regions of interest in the Papez circuit, we calculated tract-to-ROI connectivity fingerprints in people with Alzheimer’s disease (AD), mild cognitive impairment (MCI), and healthy controls, as well as macaque monkeys before and after fornix transection. We found that the hippocampal subfields’ and the anterior thalamus (ATh) connections to the fornix are dramatically degraded in AD, and (although reduced compared to controls) these connections are still present in MCI.

Our comparisons of connectivity fingerprints across groups indicate, for the first time to the best of our knowledge, that, apart from the ‘hallmark’ neurodegeneration in the hippocampus and the ATh (as landmark previous studies have shown -see *Introduction*), it is also these regions’ white matter connectivity patterns that can distinguish people with AD from people with MCI or healthy controls.

The fornix contributes key roles to memory and cognition by providing the mechanistic link between the extended hippocampal system, anterior thalamus (ATh) and mamillary bodies, and by providing the means for several neuromodulators -in particular acetylcholine-to the hippocampal formation (13, 14). As such, it is unsurprising that a compromised fornix relates to cognitive decline, as indicated by our Fractional Anisotropy findings and by several previous studies. Because of a compromised fornix in AD and MCI, it has been proposed (28) that alternative pathways may contribute to episodic memory performance, possibly via other medial temporal lobe tracts, such as the cingulum bundle or the uncinate fasciculus. Our results indicate that hippocampal connections to the cingulum bundle (CBT) remained unaffected in AD and MCI, suggesting that it is the hippocampus-fornix pathway that primarily relates to AD. We, rather, observed increased connectivity to frontal white matter tracts in the AD group (and to a lesser extent in MCI) compared to controls, that is, between the hippocampal subfields and the IFO tract, and between the ATh and the ATR tract. On the other hand, hippocampal-IFO and ATh-ATR connections were degraded in our fornix-transected animals, suggesting that a viable fornix is likely essential for the increased connections observed in the human patients. Nevertheless, we should underlie a potential limitation on the interpretation of connectivity fingerprints here. Given that these are normalised fingerprints (i.e., each fingerprint should sum to 1), it is likely that if there is a reduction in connectivity for any given tract, there will be an overall increase across other tracts (following normalisation). Hence, the possibility that the increases we observed reflect the normalisation rather than a true increase in connectivity cannot be excluded. Yet, in a recent clinical study, Ríos et al. (36) sought to identify fiber tracts associating with optimal fornix DBS outcomes, and employed a fiber filtering approach based on normative whole-brain connectivity data. They found that the fornix, and tracts that correspond to thalamic and orbitofrontal projections, accounted for optimal improvement in the training cohort that received fiber filtering. These findings accord with our finding of elevated ATh-ATR connectivity in AD, suggesting it is not simply normalisation-related. Rather, we suggest it relates to the enhanced recruitment of new neural resources as a compensatory mechanism for cognitive decline and dysfunction in other parts of the brain. (This suggestion is also informed by the increased FA we found for the AD group in the IFO tract -see *Results*). Compensatory increased task-based brain activity and increased connectivity have been reported previously in ageing (59–61) and in neurodegenerative disease (62–64). However, an established definition of compensation in neurodegeneration is missing and future research should shed light on the potential underlying mechanisms. We speculate that these may include white matter and myelin plasticity mechanisms (65, 66).

We have additionally examined structural connectivity across our ROIs, and found that the connections between all hippocampal subfields and the anterior thalamus (ATh) are strongly degraded in both hemispheres in people with AD compared to healthy controls. We also found significant decreases between the ATh and CA32/CA4/subiculum in MCI compared to controls, indicating that these connections are likely compromised early on in cognitive decline. These changes in MCI may also suggest that the ATh is critical in the transition from MCI to AD, as proposed by a recent study (67) that identified degenerated glutamatergic axon terminals connecting to neurons in the ATh. Nevertheless, our finding of reduced left ATh-CA32 connectivity in AD compared to MCI suggests that this particular connection is disrupted not before AD clinical manifestation. Considering the retrosplenial cortex (RSC), we found reduced connectivity between the RSC regions and the thalamus in people with AD and monkeys with fornix transection. Strong RSC-hippocampal CA3 degradation was also found in the post-op monkeys (but without statistical significance in the human AD group). Together, these results highlight that key mnemonic structures link through the RSC, supporting previous findings in animals (9–11, 57) and confirm the RSC-thalamic route also in the human brain. This provides support to the proposal of the RSC as an attractive candidate for the development of novel therapeutics and interventions to slow neural degeneration (9). Our fingerprint divergence results across species further indicate that the RSC’s white matter connectivity patterns resemble between humans with AD and monkeys with fornix transection. Finally, unlike the AD patients, connectivity between RSC area 30 and hippocampal/subicular subfields was increased in MCI compared to controls (although without surviving multiple comparisons corrections). This might suggest that an RSC-hippocampal route contributes to mnemonic functions still preserved in MCI, but the RSC’s engagement is no longer possible in AD and/or in cases of fornix damage.

## Materials and Methods

### Subjects

We used diffusion-weighted (dMRI) and structural T1-weighted MRI data from 20 Alzheimer’s Disease (AD) patients (mean age=74.4, 11 females), 20 people diagnosed with mild cognitive impairment (MCI) (mean age=73.8, 11 females), and 20 healthy controls (mean age=74.7, 11 females). All participants were selected from the publicly-available Alzheimer’s Disease Neuroimaging Initiative (ADNI https://adni.loni.usc.edu/) dataset, at random, ensuring that participants across groups were age- and sex-matched.

We also used a cohort of 8 male rhesus macaque monkeys (*Macaca mulatta)* aged 5 years at the beginning of the study. Four monkeys (“fornix group”) were housed together and trained (68) on the touchscreen object-in-place discrimination task (21). Subsequently, three monkeys from this group underwent neurosurgery under general anaesthesia to receive a selective bilateral fornix transection. As detailed in a previous study from our group using these monkeys (69), the fornix-transected animals, when compared to their own preoperative sessions, were significantly impaired in learning new visuospatial discriminations in the object-in-place task. The remaining four monkeys (“control group”) were housed together in a separate group, trained on a passive fixation task, received a head-post implanted onto the skull under general anaesthesia, but otherwise remained neurologically intact (70).

T1-weighted and diffusion-weighted MRI data were collected from each monkey at different time points during their individual experiments, under general anaesthesia. Scans took place at around the same time for the two groups, and the time difference between pre-op and post-op scans were equal for the two groups (∼2 years). All monkey experimental procedures were performed in compliance with the United Kingdom Animals (Scientific Procedures) Act of 1986. A Project License was reviewed by the University of Oxford Animal Care and Ethical Review Committee, the Animals in Science Committee, and the Home Office (UK) who approved and licensed all procedures. The housing and husbandry compiled with the ARRIVE guidelines of the European Directive (2010/63/EU) for the care and use of laboratory animals.

### Imaging data acquisition and pre-processing

Whole brain magnetic resonance imaging (MRI) scans of the participants in the ADNI dataset were collected using 3T GE scanners at several acquisition sites in North America. Imaging protocols can be found in https://adni.loni.usc.edu/data-samples/adni-data/neuroimaging/mri/mri-scanner-protocols/. Briefly, for the diffusion MRI data: voxel size=2.7 mm isotropic, 41 diffusion-weighted images (b value = 1,000 s/mm^2^), TR=9 s.

Whole-brain monkey MRI data were collected using a horizontal 3T scanner and a custom-made four-channel phased-array receiver coil, together with a radial transmission coil (Wind-miller Kolster Scientific). The anaesthetised monkeys were placed in the scanner in a feet first prone position with their head secured in an MRI-compatible stereotaxic frame (Crist Instrument). For details on the induction and maintenance of the anaesthesia, see (69). For dMRI data acquisition, we used an echo planar imaging (EPI) sequence with the following imaging parameters: voxel size=1 mm isotropic, b values=0 and 1,000 s/mm^2^, 60 isotropically distributed diffusion-encoding directions, TR=8.3 s, and TE=102 ms, with alternating phase-encoding directions (anterior-posterior, posterior-anterior). We collected six averages (all 60 diffusion-encoding directions and 11 b0 images in each average) in two alternating phase-encoding directions within a single diffusion-weighted scan session.

Monkey dMRI data were corrected for susceptibility field distortions along the phase-encoding direction, using FSL’s ‘Top-Up’ image distortion correction tool. Data were skull-stripped before fitting the diffusion tensor model, a particular method widely used to estimate microstructural properties. Human dMRI data were corrected for eddy current-induced distortions and subject movements using FSL’s ‘EDDY’ tool, before skull-stripping, or fitting the diffusion tensor model. Structural data were pre-processed using Freesurfer’s ‘recon-all’ pipeline. For each individual subject’s dMRI session, in each species, we also ran BEDPOSTX, an FSL tool that uses Bayesian techniques to estimate fiber orientations for up to three fiber populations per voxel (71), which were subsequently used to inform tractography (see below).

### White matter tracts of interest

We reconstructed a network of WM tracts (full list and abbreviations in Fig. 1) based on previous literature having shown that these tracts are implicated in memory and higher cognition, or exhibit diffusion MRI-derived degeneration in AD and/or MCI (25, 72–79).

Tracts were reconstructed on each individual human and macaque brain using “XTRACT” cross-species tractography (39). XTRACT uses the crossing fiber fitted data (obtained from BEDPOSTX -see above) and performs probabilistic tractography (using FSL’s probabilistic tractography algorithm PROBTRACKX (80)) to generate tracts in standard space (MNI in humans, F99 in macaques). XTRACT’s reconstruction protocols are standardised and validated, employing carefully-selected seed, target and exclusion masks to guide tractography for each tract in each species. Resulting tracts are quite robust with a consistent layout across subjects even in small sample sizes (39).

We then extracted tracts’ Fractional Anisotropy (FA) summary statistics, using “XTRACT_stats”. The FA values were analysed in R (version 4.3.3) using linear mixed-effects models implemented using the *lme4* package. As the data included multiple measurements (for the multiple tracts) from each subject, subject ID was included as a random effect in all models. Effects of subject group, tract and laterality were evaluated by comparing models with and without these factors included as fixed effects. Model comparisons used log likelihood ratio tests, reported by the chi-square and associated p-values. Post-hoc comparisons between the control, AD and MCI groups were performed with functions from the *emmeans* package, using correction for multiple comparisons according to the Dunnet method. Visual inspection of residual plots was used to detect notable deviations from normality or heteroskedasticity. Where the residual plots showed strong deviations, the models were tested for sensitivity to outliers by repeating the analysis after removing data points associated with residuals more than 2 standard deviations from the mean.

### Regions of Interest (ROIs)

In the tract-to-ROI and ROI-to-ROI analyses, we used brain regions identified as key contributors to episodic memory and established correlates of cognitive decline in humans and monkeys. Specifically, we included the hippocampal formation (hippocampal subfields: CA1, CA3/CA2, CA4 and subicular complex regions: parasubiculum, presubiculum, subiculum (4, 81, 82)), anterior thalamic nuclei (ATh) (2, 6–8), and retrosplenial cortex (regions BA 29 and 30) (9–11). Additionally, we included the mediodorsal thalamus (MD), which is adjacent to the ATh and also part of the limbic thalamus due to its reciprocal connectivity with the prefrontal cortex and inputs from medial temporal lobe structures. The MD is damaged in diencephalic amnesias along with the ATh (83).

Human thalamic nuclei, hippocampal and subicular ROIs were segmented on individual brains using each participant’s T1-weighted image, employing the methods described in (84, 85). Human CA2 and CA3 subfields are combined (85) and abbreviated as ‘CA32’, while, for the ATh, we used the anteroventral thalamus (‘AV’ in Iglesias et al. (84)) that includes the anterior medial and anterior dorsal nuclei (84). The retrosplenial cortex areas were segmented from the Harvard-Oxford cortical atlas in MNI space before co-registering to individual brains’ native space using non-linear transformation tools. In macaques, thalamic, hippocampal and subicular ROIs were delineated according to the parcellations in Saleem et al. subcortical atlas (86). We included the anteroventral (AVN) and the anteromedial (AMN) nuclei of the thalamus that together encompass the ATh. The cortical ROIs were delineated according to Van Essen et al. cortical atlas (87). For the full list of ROIs and their abbreviations, and for brain representations of core subcortical ROIs, see Fig. 6. For brain representations of the retrosplenial ROIs, see Fig. 7.

### Tract-to-ROI connectivity fingerprints and calculation of divergence

We calculated connectivity fingerprints (38–40) between our volumetric ROIs and our tracts of interest, using “XTRACT_blueprint” (https://fsl.fmrib.ox.ac.uk/fsl/docs/#/diffusion/xtract). For each brain, in each species, we ran tractography from each ROI towards whole-brain white matter in native space (then transformed to standard space -MNI in humans; F99 in macaques). This produced a connectivity matrix [ROI x white matter]. Tracts of interest (generated by XTRACT -see above) were then converted to vectors and stacked to create a second matrix [white matter x tracts]. Finally, the two matrices were multiplied to produce the connectivity blueprint, that is a matrix [ROI x tracts] (Fig. 8). We averaged the connectivity blueprint across voxels for a given ROI, then averaged across subjects belonging to the same group, to get the connectivity fingerprint/pattern of each ROI (Fig. 1-3).

**Fig. 8.**
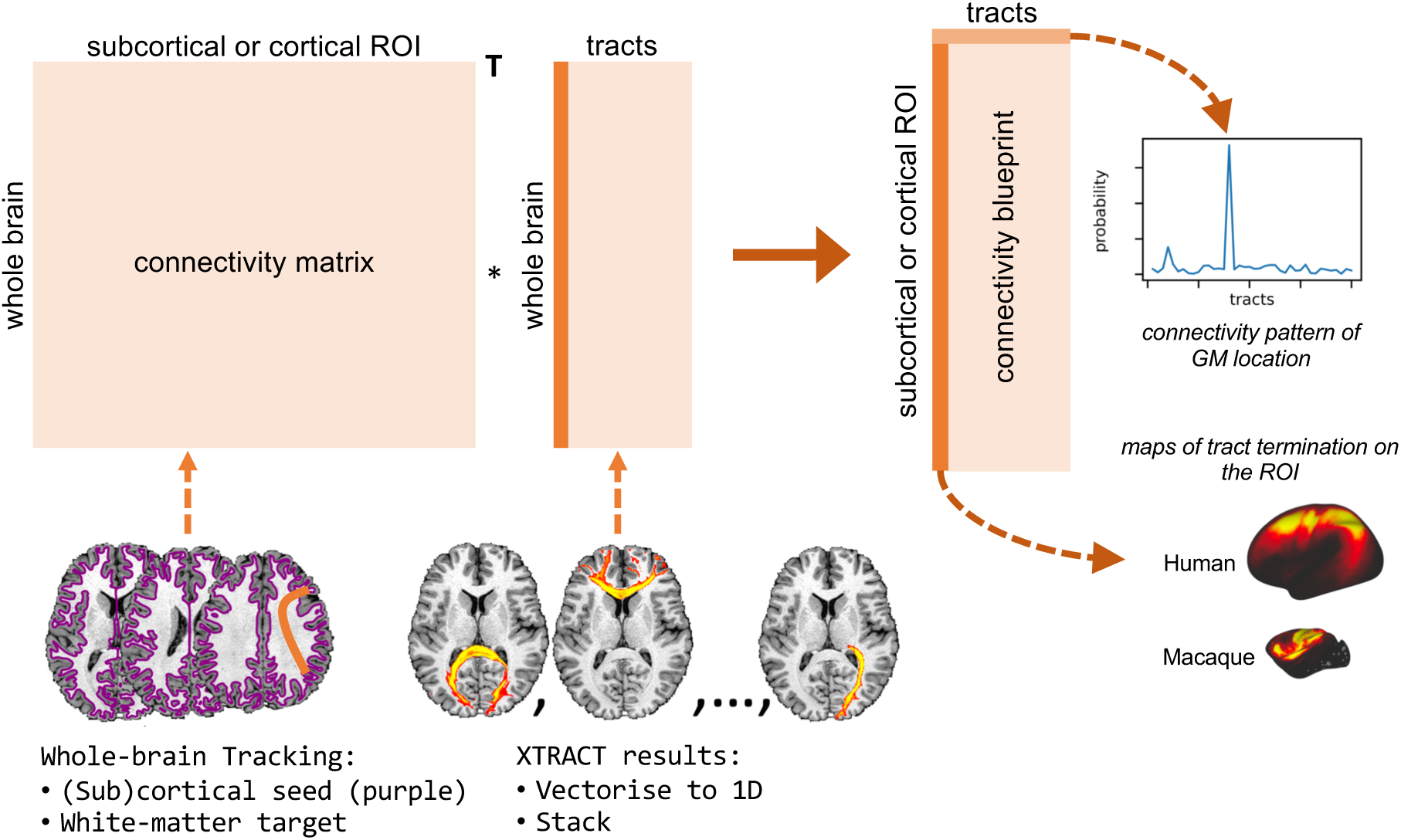
Schematic of the connectivity fingerprint generation pipeline. Columns in the connectivity blueprint matrix (right) provide maps of (subcortical or cortical) ROIs where tracts terminate, while rows consist of a ROIs’ connectivity patterns showing how each ROI connects to the tracts. Adapted from Warrington et al. (40).

We then created group-averaged connectivity fingerprints for each ROI, in each species. Group-averaged connectivity fingerprints were fed to an adapted version of “XTRACT_divergence”, which compares fingerprints across groups (or across brains / species (38)). We adopted Kullback-Leibler (KL) divergence, a statistical metric that compares probability distributions, to demonstrate, in each ROI, the value of divergence between any given voxel in one group (or species) and the ROI’s mean fingerprint in another group (or species). We quantified the KL divergence maps’ voxels values and plotted divergence distributions as histograms (Fig. 4, Fig. 7).

### ROI-to-ROI connectivity

We used PROBTRACKX (80), which produces sample streamlines starting from any seed (here defined as every voxel in an entire volumetric region). The algorithm then uses the sample streamlines to estimate the number of streamlines that visited each voxel.

We have selected 11 ROIs (see *Regions of Interest*), examined for both hemispheres. Hence, streamlines were seeded from each ROI mask, with PROBTRACKX estimating the number of connecting streamlines ROI x ROI, ending up with a 22×22 connectivity matrix per MRI session for humans, and a 24×24 matrix for NHPs (since ATh was subdivided in AVN and AMN). Each row in the averaged matrices in Fig. 6 quantifies the number of streamlines that seeded in a given ROI and reached the other ROIs. To account for the inter-subject, inter-session, and inter-species variability in the number of generated streamlines, we normalised the number of streamlines within each session by dividing each ROI-ROI pair streamline number by the total number of streamlines generated from each seed ROI.

To evaluate group differences in structural connectivity, we calculated, for every ROI-ROI pair, the difference in the average (across subjects in a given group) number of streamlines between two groups (see also 88). For example, the [AD – Controls] test assessed the connectivity changes in AD compared to the control group. The statistical significance of these differences was assessed by permutation tests, where we randomly shuffled the labels of subjects’ groups. For each permutation, the absolute value of the difference between (the shuffled) labels was calculated. ROI-ROI pairs for which the real absolute difference was larger than the 95th percentile (one-tailed) of the null distribution (calculated after shuffling) are indicated with one asterisk in Fig. 6. Two asterisks indicate ROI-ROI pairs for which the real absolute difference was larger than the 95th percentile, FDR-corrected for the multiple comparisons across the ROI-ROI pairs (89).

### Fornix transection

Surgical procedures, histology and the assessment of fornix transections in our monkeys have been detailed previously (69, 90). Briefly, three monkeys in the ‘fornix group’ underwent a bilateral transection of the fornix in the region above either the intraventricular foramen or the anterior thalamus. A small hole was made at the midline in the corpus callosum extending to ∼8-10 mm in length from above the intraventricular foramen to above the mediodorsal thalamus. One monkey received unilateral fornix transection, and this animal’s post-op scan session was not included in the postoperative data analyses.

## Author contributions

V.P. conceptualised the study; V.P., S.N.S, A.S.M. designed research; V.P., A.S.M. performed research; V.P., S.W., E.P., J.D.B., S.N.S, A.S.M. contributed analytic tools; V.P. wrote the paper; V.P., S.W., E.P., J.D.B., S.N.S, A.S.M. revised the manuscript.

## Acknowledgements

V.P. is supported by a Medical Research Foundation grant (MRF-RGM-HR-23-101). S.W. and S.N.S. are supported by an ERC Consolidator grant (101000969). The monkey research was supported by funding from the Medical Research Council UK (G0800329) and Wellcome Trust WT (110157/Z/15/Z) to A.S.M. We thank Oxford University Biomedical Services animal technicians and veterinarians for support with anaesthesia, post procedure recovery, and monitoring.

Data collection and sharing for this project was funded by the Alzheimer’s Disease Neuroimaging Initiative (ADNI) (National Institutes of Health Grant U01 AG024904) and DOD ADNI (Department of Defense award number W81XWH-12-2-0012). ADNI is funded by the National Institute on Aging, the National Institute of Biomedical Imaging and Bioengineering, and through generous contributions from the following: AbbVie, Alzheimer’s Association; Alzheimer’s Drug Discovery Foundation; Araclon Biotech; BioClinica, Inc.; Biogen; Bristol-Myers Squibb Company; CereSpir, Inc.; Cogstate; Eisai Inc.; Elan Pharmaceuticals, Inc.; Eli Lilly and Company; EuroImmun; F. Hoffmann-La Roche Ltd and its affiliated company Genentech, Inc.; Fujirebio; GE Healthcare; IXICO Ltd.; Janssen Alzheimer Immunotherapy Research & Development, LLC.; Johnson & Johnson Pharmaceutical Research & Development LLC.; Lumosity; Lundbeck; Merck & Co., Inc.; Meso Scale Diagnostics, LLC.; NeuroRx Research; Neurotrack Technologies; Novartis Pharmaceuticals Corporation; Pfizer Inc.; Piramal Imaging; Servier; Takeda Pharmaceutical Company; and Transition Therapeutics. The Canadian Institutes of Health Research is providing funds to support ADNI clinical sites in Canada. Private sector contributions are facilitated by the Foundation for the National Institutes of Health (www.fnih.org). The grantee organization is the Northern California Institute for Research and Education, and the study is coordinated by the Alzheimer’s Therapeutic Research Institute at the University of Southern California. ADNI data are disseminated by the Laboratory for Neuro Imaging at the University of Southern California.

---

**Supplementary Fig. 1.**
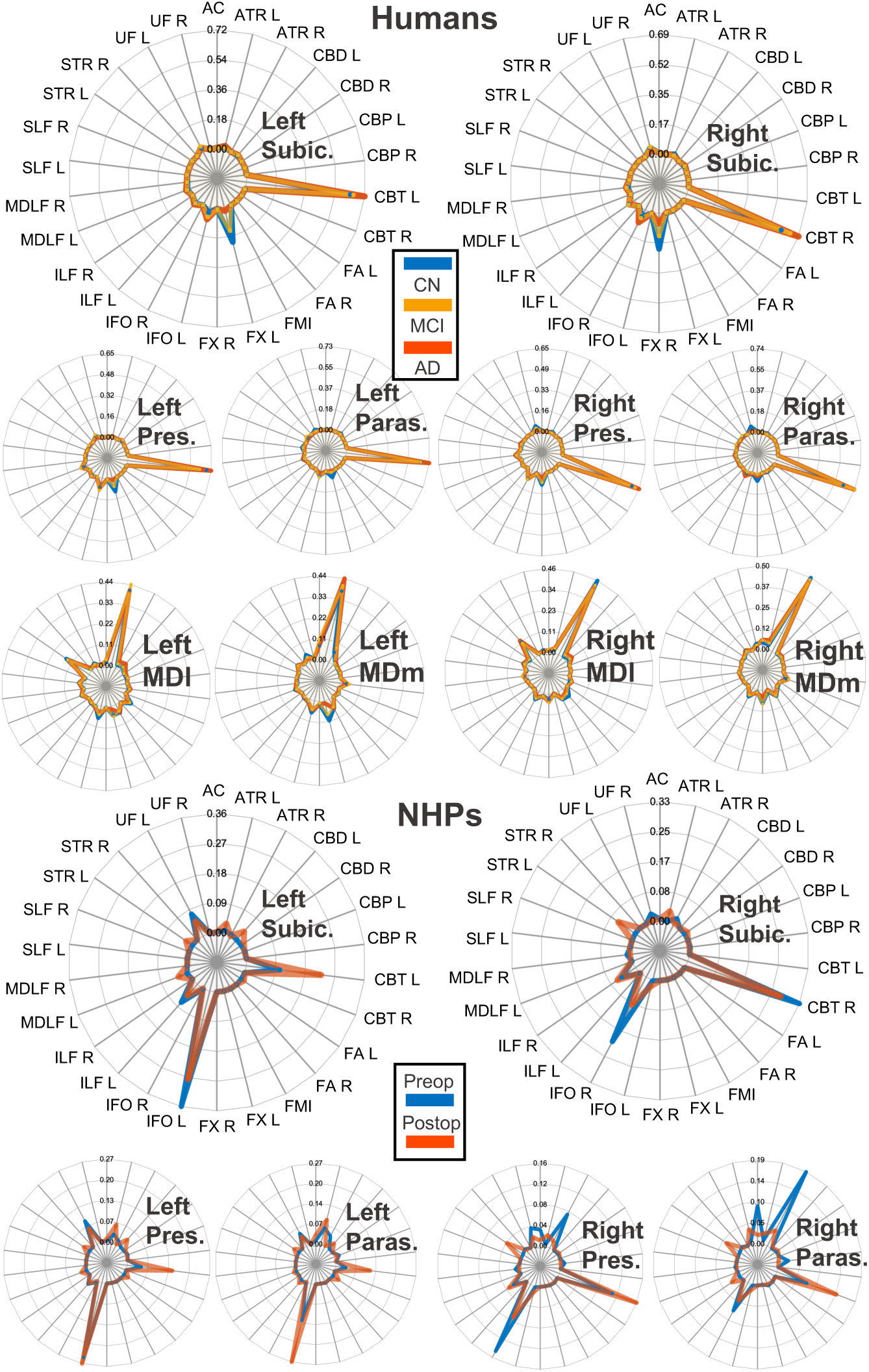
Connectivity fingerprints between subicular complex regions and white matter tracts, in the left and right hemisphere. Top panel shows human data for healthy controls (CN), mild cognitive impairment (MCI) and Alzheimer’s disease (AD) patients. A row with the fingerprints between the MD thalamic nuclei and white matter tracts also shown. Bottom panel shows the non-human primate (NHP) data before (pre-op) and after (post-op) the fornix transection operations. Given that the fornix was transected in the post-op group, connectivity fingerprints to the fornix were not calculated. ROI abbreviations: subic: subiculum; pres: presubiculum; paras: parasubiculum; MDl: mediodorsal thalamus lateral (parvocellular); MDm: mediodorsal thalamus medial (magnocellular). Tracts abbreviations: fx: Fornix; atr: Anterior Thalamic Radiation; cbd: Cingulum subsection: dorsal; cbp: Cingulum subsection: perigenual; cbt: Cingulum subsection: temporal; fa: Frontal Aslant; ifo: Inferior Fronto-Occipital Fasciculus; ilf: Inferior Longitudinal Fasciculus; mdlf: Middle Longitudinal Fasciculus; slf1: Superior Longitudinal Fasciculus 1; str: Superior Thalamic Radiation; uf: Uncinate Fasciculus; ac: Anterior Commissure; fmi: Forceps Minor. L: left hemisphere; R: right hemisphere.

## Notes

https://adni.loni.usc.edu/

### Competing Interest Statement

The authors have declared no competing interest.

## References

1. Anonymous, 2024 Alzheimer’s disease facts and figures. Alzheimers Dement 20, 3708–3821 (2024).

2. J. P. Aggleton, A. Pralus, A. J. Nelson, M. Hornberger, Thalamic pathology and memory loss in early Alzheimer’s disease: moving the focus from the medial temporal lobe to Papez circuit. Brain 139, 1877–1890 (2016).

3. J. W. Papez, A proposed mechanism of emotion. 1937. J Neuropsychiatry Clin Neurosci 7, 103–112 (1995).

4. H. Braak, E. Braak, Neuropathological stageing of Alzheimer-related changes. Acta Neuropathol 82, 239–259 (1991).

5. W. J. Henneman et al., Hippocampal atrophy rates in Alzheimer disease: added value over whole brain volume measures. Neurology 72, 999–1007 (2009).

6. J. P. Aggleton, S. M. O’Mara, The anterior thalamic nuclei: core components of a tripartite episodic memory system. Nat Rev Neurosci 23, 505–516 (2022).

7. H. Braak, E. Braak, Alzheimer’s disease affects limbic nuclei of the thalamus. Acta Neuropathol 81, 261–268 (1991).

8. J. M. Biesbroek, M. G. Verhagen, S. van der Stigchel, G. J. Biessels, When the central integrator disintegrates: A review of the role of the thalamus in cognition and dementia. Alzheimers Dement 20, 2209–2222 (2024).

9. S. Trask, D. I. Fournier, Examining a role for the retrosplenial cortex in age-related memory impairment. Neurobiol Learn Mem 189, 107601 (2022).

10. E. Valenstein et al., Retrosplenial amnesia. Brain 110 (Pt 6), 1631–1646 (1987).

11. S. D. Vann, J. P. Aggleton, E. A. Maguire, What does the retrosplenial cortex do? Nat Rev Neurosci 10, 792–802 (2009).

12. Y. Chen et al., Abnormal white matter changes in Alzheimer’s disease based on diffusion tensor imaging: A systematic review. Ageing Res Rev 87, 101911 (2023).

13. C. E. Poletti, G. Creswell, Fornix system efferent projections in the squirrel monkey: an experimental degeneration study. J Comp Neurol 175, 101–128 (1977).

14. R. C. Saunders, J. P. Aggleton, Origin and topography of fibers contributing to the fornix in macaque monkeys. Hippocampus 17, 396–411 (2007).

15. R. C. Petersen et al., Current concepts in mild cognitive impairment. Arch Neurol 58, 1985–1992 (2001).

16. R. A. Sperling et al., Toward defining the preclinical stages of Alzheimer’s disease: recommendations from the National Institute on Aging-Alzheimer’s Association workgroups on diagnostic guidelines for Alzheimer’s disease. Alzheimers Dement 7, 280–292 (2011).

17. E. Fletcher et al., Loss of fornix white matter volume as a predictor of cognitive impairment in cognitively normal elderly individuals. JAMA Neurol 70, 1389–1395 (2013).

18. M. M. Mielke et al., Fornix integrity and hippocampal volume predict memory decline and progression to Alzheimer’s disease. Alzheimers Dement 8, 105–113 (2012).

19. J. P. Aggleton et al., Differential cognitive effects of colloid cysts in the third ventricle that spare or compromise the fornix. Brain 123 ( Pt 4), 800–815 (2000).

20. M. D’Esposito, M. Verfaellie, M. P. Alexander, D. I. Katz, Amnesia following traumatic bilateral fornix transection. Neurology 45, 1546–1550 (1995).

21. D. Gaffan, Scene-specific memory for objects: a model of episodic memory impairment in monkeys with fornix transection. J Cogn Neurosci 6, 305–320 (1994).

22. A. Poreh et al., Anterograde and retrograde amnesia in a person with bilateral fornix lesions following removal of a colloid cyst. Neuropsychologia 44, 2241–2248 (2006).

23. D. Tsivilis et al., A disproportionate role for the fornix and mammillary bodies in recall versus recognition memory. Nat Neurosci 11, 834–842 (2008).

24. C. R. Wilson, D. P. Charles, M. J. Buckley, D. Gaffan, Fornix transection impairs learning of randomly changing object discriminations. J Neurosci 27, 12868–12873 (2007).

25. M. M. Mielke et al., Regionally-specific diffusion tensor imaging in mild cognitive impairment and Alzheimer’s disease. Neuroimage 46, 47–55 (2009).

26. R. D. Perea et al., Connectome-derived diffusion characteristics of the fornix in Alzheimer’s disease. Neuroimage Clin 19, 331–342 (2018).

27. I. Shaikh et al., Diffusion tensor tractography of the fornix in cerebral amyloid angiopathy, mild cognitive impairment and Alzheimer’s disease. Neuroimage Clin 34, 103002 (2022).

28. C. Metzler-Baddeley et al., Temporal association tracts and the breakdown of episodic memory in mild cognitive impairment. Neurology 79, 2233–2240 (2012).

29. L. Zhuang et al., Abnormalities of the fornix in mild cognitive impairment are related to episodic memory loss. J Alzheimers Dis 29, 629–639 (2012).

30. R. Li, C. Zhang, Y. Rao, T. F. Yuan, Deep brain stimulation of fornix for memory improvement in Alzheimer’s disease: A critical review. Ageing Res Rev 79, 101668 (2022).

31. A. M. Lozano et al., A Phase II Study of Fornix Deep Brain Stimulation in Mild Alzheimer’s Disease. J Alzheimers Dis 54, 777–787 (2016).

32. C. Hamani et al., Memory enhancement induced by hypothalamic/fornix deep brain stimulation. Ann Neurol 63, 119–123 (2008).

33. M. Bittlinger, S. Müller, Opening the debate on deep brain stimulation for Alzheimer disease - a critical evaluation of rationale, shortcomings, and ethical justification. BMC Med Ethics 19, 41 (2018).

34. Z. Liu, K. Shu, Y. Geng, C. Cai, H. Kang, Deep brain stimulation of fornix in Alzheimer’s disease: From basic research to clinical practice. Eur J Clin Invest 53, e13995 (2023).

35. S. Senova, A. Fomenko, E. Gondard, A. M. Lozano, Anatomy and function of the fornix in the context of its potential as a therapeutic target. J Neurol Neurosurg Psychiatry 91, 547–559 (2020).

36. A. S. Ríos et al., Optimal deep brain stimulation sites and networks for stimulation of the fornix in Alzheimer’s disease. Nat Commun 13, 7707 (2022).

37. G. S. Smith et al., Increased cerebral metabolism after 1 year of deep brain stimulation in Alzheimer disease. Arch Neurol 69, 1141–1148 (2012).

38. R. B. Mars et al., Whole brain comparative anatomy using connectivity blueprints. Elife 7 (2018).

39. S. Warrington et al., XTRACT - Standardised protocols for automated tractography in the human and macaque brain. Neuroimage 217, 116923 (2020).

40. S. Warrington et al., Concurrent mapping of brain ontogeny and phylogeny within a common space: Standardized tractography and applications. Sci Adv 8, eabq2022 (2022).

41. M. Lacalle-Aurioles, Y. Iturria-Medina, Fornix degeneration in risk factors of Alzheimer’s disease, possible trigger of cognitive decline. Cereb Circ Cogn Behav 4, 100158 (2023).

42. S. Assimopoulos et al., Generalising XTRACT tractography protocols across common macaque brain templates. Brain Struct Funct 229, 1873–1888 (2024).

43. M. A. Ferguson et al., A human memory circuit derived from brain lesions causing amnesia. Nat Commun 10, 3497 (2019).

44. N. McNaughton, S. D. Vann, Construction of complex memories via parallel distributed cortical-subcortical iterative integration. Trends Neurosci 45, 550–562 (2022).

45. H. Johansen-Berg, M. F. Rushworth, Using diffusion imaging to study human connectional anatomy. Annu Rev Neurosci 32, 75–94 (2009).

46. D. K. Jones, T. R. Knösche, R. Turner, White matter integrity, fiber count, and other fallacies: the do’s and don’ts of diffusion MRI. Neuroimage 73, 239–254 (2013).

47. L. W. Wang et al., White and gray matter integrity evaluated by MRI-DTI can serve as noninvasive and reliable indicators of structural and functional alterations in chronic neurotrauma. Sci Rep 14, 7244 (2024).

48. Z. Zhou et al., Evaluation of the diffusion MRI white matter tract integrity model using myelin histology and Monte-Carlo simulations. Neuroimage 223, 117313 (2020).

49. G. Bartzokis et al., Heterogeneous age-related breakdown of white matter structural integrity: implications for cortical “disconnection” in aging and Alzheimer’s disease. Neurobiol Aging 25, 843–851 (2004).

50. N. Geschwind, Disconnexion syndromes in animals and man. I. Brain 88, 237–294 (1965).

51. M. O’Sullivan et al., Evidence for cortical “disconnection” as a mechanism of age-related cognitive decline. Neurology 57, 632–638 (2001).

52. C. Metzler-Baddeley, D. K. Jones, B. Belaroussi, J. P. Aggleton, M. J. O’Sullivan, Frontotemporal connections in episodic memory and aging: a diffusion MRI tractography study. J Neurosci 31, 13236–13245 (2011).

53. I. J. Bennett, D. J. Huffman, C. E. Stark, Limbic Tract Integrity Contributes to Pattern Separation Performance Across the Lifespan. Cereb Cortex 25, 2988–2999 (2015).

54. L. Zhuang et al., Microstructural white matter changes in cognitively normal individuals at risk of amnestic MCI. Neurology 79, 748–754 (2012).

55. A. Pelletier et al., Structural hippocampal network alterations during healthy aging: a multi-modal MRI study. Front Aging Neurosci 5, 84 (2013).

56. J. P. Aggleton, A. J. D. Nelson, S. M. O’Mara, Time to retire the serial Papez circuit: Implications for space, memory, and attention. Neurosci Biobehav Rev 140, 104813 (2022).

57. A. S. Mitchell, R. Czajkowski, N. Zhang, K. Jeffery, A. J. D. Nelson, Retrosplenial cortex and its role in spatial cognition. Brain Neurosci Adv 2, 2398212818757098 (2018).

58. E. J. Bubb, L. Kinnavane, J. P. Aggleton, Hippocampal - diencephalic - cingulate networks for memory and emotion: An anatomical guide. Brain Neurosci Adv 1 (2017).

59. R. Cabeza et al., Maintenance, reserve and compensation: the cognitive neuroscience of healthy ageing. Nat Rev Neurosci 19, 701–710 (2018).

60. U. Lindenberger, Human cognitive aging: corriger la fortune? Science 346, 572–578 (2014).

61. A. Salami, S. Pudas, L. Nyberg, Elevated hippocampal resting-state connectivity underlies deficient neurocognitive function in aging. Proc Natl Acad Sci U S A 111, 17654–17659 (2014).

62. D. Berron, D. van Westen, R. Ossenkoppele, O. Strandberg, O. Hansson, Medial temporal lobe connectivity and its associations with cognition in early Alzheimer’s disease. Brain 143, 1233–1248 (2020).

63. S. Gregory, J. D. Long, S. J. Tabrizi, G. Rees, Measuring compensation in neurodegeneration using MRI. Curr Opin Neurol 30, 380–387 (2017).

64. L. Pasquini et al., Link between hippocampus’ raised local and eased global intrinsic connectivity in AD. Alzheimers Dement 11, 475–484 (2015).

65. C. Sampaio-Baptista, H. Johansen-Berg, White Matter Plasticity in the Adult Brain. Neuron 96, 1239–1251 (2017).

66. W. Xin, J. R. Chan, Myelin plasticity: sculpting circuits in learning and memory. Nat Rev Neurosci 21, 682–694 (2020).

67. B. Sárkány et al., Early and selective localization of tau filaments to glutamatergic subcellular domains within the human anterodorsal thalamus. Acta Neuropathol 147, 98 (2024).

68. S. Mason et al., Effective chair training methods for neuroscience research involving rhesus macaques (Macaca mulatta). J Neurosci Methods 317, 82–93 (2019).

69. V. Pelekanos et al., Corticocortical and Thalamocortical Changes in Functional Connectivity and White Matter Structural Integrity after Reward-Guided Learning of Visuospatial Discriminations in Rhesus Monkeys. J Neurosci 40, 7887–7901 (2020).

70. V. Pelekanos et al., Rapid event-related, BOLD fMRI, non-human primates (NHP): choose two out of three. Sci Rep 10, 7485 (2020).

71. T. E. Behrens, H. J. Berg, S. Jbabdi, M. F. Rushworth, M. W. Woolrich, Probabilistic diffusion tractography with multiple fibre orientations: What can we gain? Neuroimage 34, 144–155 (2007).

72. E. J. Bubb, C. Metzler-Baddeley, J. P. Aggleton, The cingulum bundle: Anatomy, function, and dysfunction. Neurosci Biobehav Rev 92, 104–127 (2018).

73. D. N. Bullock et al., A taxonomy of the brain’s white matter: twenty-one major tracts for the 21st century. Cereb Cortex 32, 4524–4548 (2022).

74. H. Huang et al., Distinctive disruption patterns of white matter tracts in Alzheimer’s disease with full diffusion tensor characterization. Neurobiol Aging 33, 2029–2045 (2012).

75. A. Madhavan et al., Characterizing White Matter Tract Degeneration in Syndromic Variants of Alzheimer’s Disease: A Diffusion Tensor Imaging Study. J Alzheimers Dis 49, 633–643 (2016).

76. R. Mito et al., Fibre-specific white matter reductions in Alzheimer’s disease and mild cognitive impairment. Brain 141, 888–902 (2018).

77. F. Palesi et al., Specific Patterns of White Matter Alterations Help Distinguishing Alzheimer’s and Vascular Dementia. Front Neurosci 12, 274 (2018).

78. R. J. Von Der Heide, L. M. Skipper, E. Klobusicky, I. R. Olson, Dissecting the uncinate fasciculus: disorders, controversies and a hypothesis. Brain 136, 1692–1707 (2013).

79. Q. Wen et al., White matter alterations in early-stage Alzheimer’s disease: A tract-specific study. Alzheimers Dement (Amst) 11, 576–587 (2019).

80. T. E. Behrens et al., Characterization and propagation of uncertainty in diffusion-weighted MR imaging. Magn Reson Med 50, 1077–1088 (2003).

81. S. L. Ding, Comparative anatomy of the prosubiculum, subiculum, presubiculum, postsubiculum, and parasubiculum in human, monkey, and rodent. J Comp Neurol 521, 4145–4162 (2013).

82. S. O’Mara, The subiculum: what it does, what it might do, and what neuroanatomy has yet to tell us. J Anat 207, 271–282 (2005).

83. G. Pergola et al., The Regulatory Role of the Human Mediodorsal Thalamus. Trends Cogn Sci 22, 1011–1025 (2018).

84. J. E. Iglesias et al., A probabilistic atlas of the human thalamic nuclei combining ex vivo MRI and histology. Neuroimage 183, 314–326 (2018).

85. J. E. Iglesias et al., A computational atlas of the hippocampal formation using ex vivo, ultra-high resolution MRI: Application to adaptive segmentation of in vivo MRI. Neuroimage 115, 117–137 (2015).

86. K. S. Saleem et al., High-resolution mapping and digital atlas of subcortical regions in the macaque monkey based on matched MAP-MRI and histology. Neuroimage 245, 118759 (2021).

87. D. C. Van Essen, M. F. Glasser, D. L. Dierker, J. Harwell, Cortical parcellations of the macaque monkey analyzed on surface-based atlases. Cereb Cortex 22, 2227–2240 (2012).

88. V. Pelekanos, E. Premereur, A. S. Mitchell, Structural Connectivity Changes After Fornix Transection in Macaques Using Probabilistic Diffusion Tractography. Adv Exp Med Biol 1423, 11–20 (2023).

89. Y. H. Benjamini, Y., Controlling the false discovery rate: a practical and powerful approach to multiple testing. J R Stat Soc B 57, 289–300 (1995).

90. A. S. Mitchell, P. G. Browning, C. R. Wilson, M. G. Baxter, D. Gaffan, Dissociable roles for cortical and subcortical structures in memory retrieval and acquisition. J Neurosci 28, 8387–8396 (2008).

